# A phased genome assembly for allele-specific analysis in *Trypanosoma brucei*

**DOI:** 10.1101/2021.04.13.439624

**Authors:** RO Cosentino, BG Brink, TN Siegel

## Abstract

Many eukaryotic organisms are diploid or even polyploid, i.e. they harbour two or more independent copies of each chromosome. Yet, to date most reference genome assemblies represent a mosaic consensus sequence in which the homologous chromosomes have been collapsed into one sequence. This procedure generates sequence artefacts and impedes analyses of allele-specific mechanisms. Here, we report the allele-specific genome assembly of the diploid unicellular protozoan parasite *Trypanosoma brucei*.

As a first step, we called variants on the allele-collapsed assembly of the *T. brucei* Lister 427 isolate using short-read error-corrected PacBio reads. We identified ~96 thousand heterozygote variants across the genome (average of 4.2 variants / kb), and observed that the variant density along the chromosomes was highly uneven. Several long (>100 kb) regions of loss-of-heterozigosity (LOH) were identified, suggesting recent recombination events between the alleles. By analysing available genomic sequencing data of multiple Lister 427 derived clones, we found that most LOH regions were conserved, except for some that were specific to clones adapted to the insect lifecycle stage. Surprisingly, we also found that some Lister 427 clones were aneuploid. We found evidence of trisomy in chromosome five (chr 5), chr 2, chr 6 and chr 7. Moreover, by analysing RNA-seq data, we showed that the transcript level is proportional to the ploidy, evidencing the lack of a general expression control at the transcript level in *T. brucei*.

As a second step, to generate an allele-specific genome assembly, we used two powerful datatypes for haplotype reconstruction: raw long reads (PacBio) and chromosome conformation (Hi-C) data. With this approach, we were able to assign 99.5% of all heterozygote variants to a specific homologous chromosome, building a 66 Mb long *T. brucei* Lister 427 allele-specific genome assembly. Hereby, we identified genes with allele-specific premature termination codons and showed that differences in allele-specific expression at the level of transcription and translation can be accurately monitored with the fully phased genome assembly.

The obtained reference-grade allele-specific genome assembly of *T. brucei* will enable the analysis of allele-specific phenomena, as well as the better understanding of recombination and evolutionary processes. Furthermore, it will serve as a standard to ‘benchmark’ much needed automatic genome assembly pipelines for highly heterozygous wild species isolates.

## Introduction

Genomes contain all genetic material of an organism, thus representing the ultimate template of heredity. Today they also lay the foundation for most molecular research. Therefore, incomplete or erroneous genome assemblies have a broad impact, constraining the spectrum of possible analyses, masking findings and delaying research.

The advent of high throughput short read sequencing technologies 15 years ago has resulted in an enormous increase of draft genomes (~1500% increase in the 2005-2012 period [1]). Yet, most of those genomes were left incomplete and highly fragmented, due to the limitation of short-read sequences, failing to resolve most repetitive regions. This scenario started to change with the more recent deployment of long-read sequencing technologies such as PacBio and Nanopore, which have dramatically improved the contiguity of genome assemblies [2, 3]. Furthermore, complementation of long-read sequencing with methods that capture chromosome conformation, like Hi-C, enables accurate scaffolding, often reaching chromosome-scale assemblies [4]. Hi-C is a high-throughput assay that measures the physical contact frequency between pairs of DNA regions on a genome-wide scale by paired-end sequencing. Even though this approach was developed to study the 3D organization of genomes, it was quickly observed that the frequency of interaction between any given pair of sequences was highly dependent on the distance between them in the linear chromosome. Thus, this information could be used to order chromosome pieces in genome assembly projects. Since then, the usage of Hi-C data has shown to be one of the most powerful scaffolding approaches.

Ploidy refers to the number of complete chromosome sets in a cell. Many eukaryotic organisms have two (diploid) or more than two (polyploid) sets of chromosomes. These chromosome sets are often not exactly identical to each other, and gene alleles can be differently regulated in the context of the alternative homologous chromosomes. In fact, an increasing body of evidence shows that allelic biased gene expression is an important factor in normal cell development and in the emergence of disease. For example, a recent study showed that monoallelic gene expression is prevalent in human brain cells, possibly playing a role in the complexity and diversity of brain cell functions, which may also explain the variable penetrance of neuronal disorders-related mutations [5].

Yet, to date most reference genome assemblies of diploid (and polyploid) organisms represent a mosaic consensus sequence for which the different alleles have been collapsed into one sequence per chromosome, not corresponding to any of the true chromosomal alleles. This procedure introduces sequence artefacts in the assembly, leading to annotation and analysis errors [6, 7]. It, furthermore, disregards the information of true allelic variants and the association between them, essential for the analysis of allele-specific mechanisms [8, 9], the identification of recombination events [10] or evolutionary studies [11].

To recover the haplotypes from a collapsed assembly, typically a two-step procedure is employed [12]. First, the positions and the possible alleles for each of the heterozygous sites are determined. Second, the co-occurrence of alleles in neighbouring sites along the chromosomes is defined. The first step can be achieved with regular short-read sequencing data and established variant callers, like FreeBayes [13] or GATK [14]. However, the main reason for the absence of allele-specific genome assemblies was the lack of sequencing technologies and dedicated software enabling the second step, the linkage of neighbouring variant sites, in an accurate manner. Albeit, the increasing availability of long read sequencing, such as PacBio or Nanopore, and short-read sequencing data capable of connecting distal regions from the same chromosomal allele, like those derived from Hi-C experiments, together with computational tools able of integrating information from different sequencing technologies to link variants, such as HapCUT2 [15], are starting to make the generation of haplotype specific genome assemblies possible.

*Trypanosoma brucei* is a diploid eukaryotic parasite, the causal agent of sleeping sickness in humans and nagana in mammals, and a model organism for the study of antigenic variation. More than 15 years ago, the publication of the first genome assembly from the *T. brucei* TREU 927 isolate [16] was a major breakthrough and an invaluable resource for the research community. However, to generate the assembly, both haplotypes were collapsed into one mosaic sequence. Recently, combining PacBio sequencing and Hi-C data, we assembled the genome of the *T. brucei* Lister 427 isolate (Tb427v9) [17], the isolate most widely used in laboratory settings. While the Tb427v9 assembly contained phased subtelomeric regions, the central part of the chromosomes encoding most of the transcriptome remained collapsed, disregarding the information of allelic variants and the association between them.

To address these limitations, we aimed to reveal the allelic variation in the *T. brucei* Lister 427 isolate genome and to reconstruct the haplotypes, generating an allele-specific genome assembly. The newly assembled phased genome will open opportunities to the trypanosome community to explore a new level of gene regulation, namely the allele specific expression.

## Results

### Outline of the genome phasing strategy

In order to generate an allele-specific genome assembly of the *T. brucei* Lister 427 isolate, we established the following strategy: a) error-corrected PacBio reads were mapped to our previously published, collapsed genome assembly (Tb427v9) [17], b) variant positions were identified and those variants suggesting errors in the assembly were corrected, c) the distribution of variants across the chromosomes was analysed, as well as the ploidy based on different gDNA-seq datasets from Lister 427 clones, d) using Hi-C data and raw PacBio reads, the variants were linked, to generate an allele-specific assembly (outlined in **Figure 1**).

**Figure 1.**
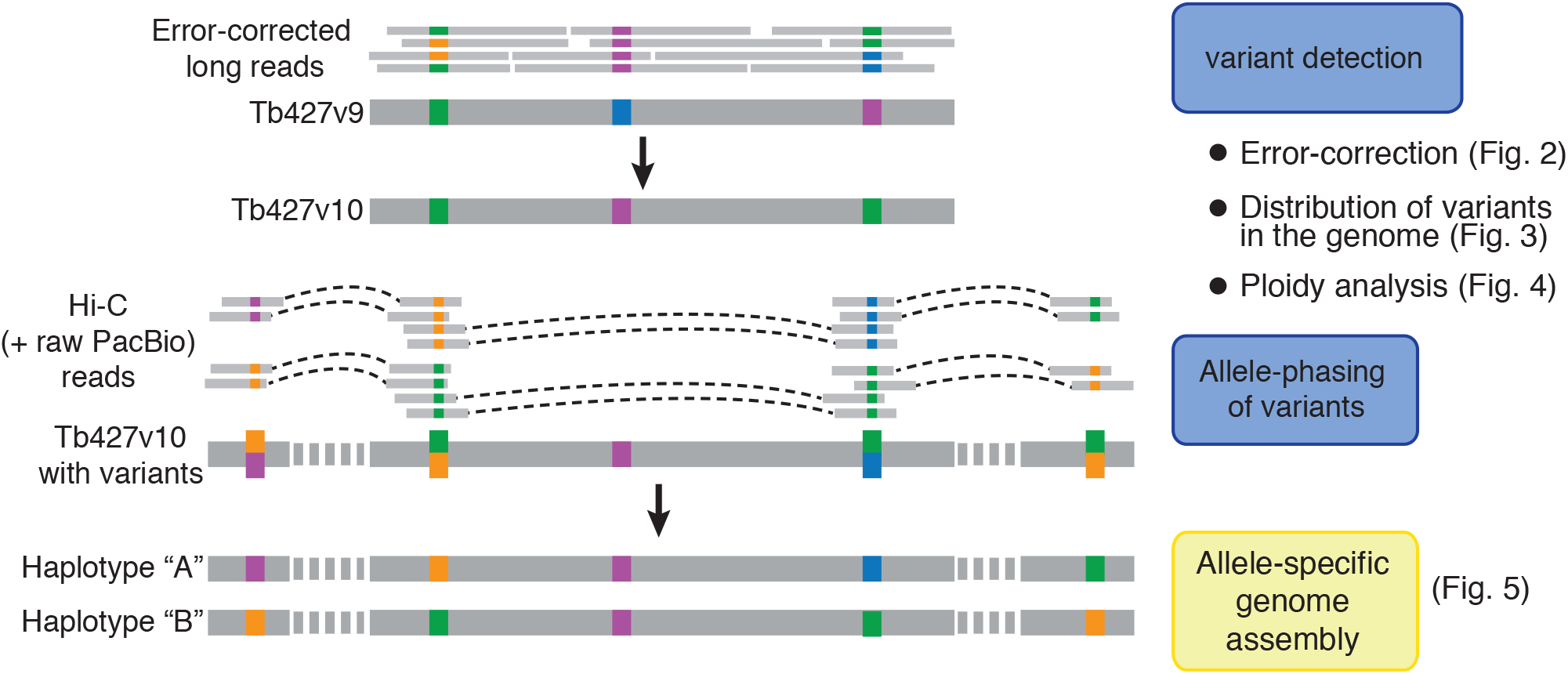
Haplotype phasing procedure. Short-read error-corrected PacBio reads were used to identify variants and fix errors in the *T. brucei* Lister 427 Tb427v9 genome assembly. Then, the distribution of heterozygous variants along the genome and the ploidy of *T. brucei* Lister 427 clones was analysed. Finally, Hi-C data and raw PacBio reads were used to link variants and reconstruct the haplotypes of the homologous chromosomes.

### Error-correction step improves accuracy of protein coding gene annotation

Genome assemblies based on long-read technologies are usually more contiguous than those based on short-reads, but at the nucleotide-level they are less accurate [18]. Moreover, the predominant errors that these technologies introduce are insertions or deletions (INDELs) [19], which, if located within an open-reading-frame, often lead to artificial truncation and the incorrect annotation of protein-coding genes as pseudogenes [20].

Thus, in order to increase the accuracy of the Lister 427 genome assembly, we performed an error-correction step before generating an allele-specific genome assembly. To this end, we first produced highly accurate long reads by correcting raw PacBio reads with Illumina short-reads [21]. Then, we mapped the error-corrected PacBio reads to the collapsed Lister 427 genome assembly v9 (Tb427v9), identified and classified variant positions. The genome sequence was corrected in the positions where the mapped reads suggested errors and the impact of the correction in terms of protein-coding gene annotation was assessed (**Figure 2a**). Using the error-corrected PacBio reads we identified more than 98,000 variant positions in the diploid ‘core’ of the Tb427v9 genome assembly. Even though most of these variants were of the ‘expected type’, i.e. heterozygotes between the allele present in the assembly and an alternative allele (variants nomenclated as “Ref/Alt1”), around 7,000 variants suggested errors in the genome assembly (**Figure 2b**). Those ‘error-suggesting variants’ were either homozygous for an alternative allele (nomenclated as “Alt1/Alt1”) or heterozygous between two alternative alleles (nomenclated as “Alt1/Alt2”). As expected by the error bias of long-read-based assemblies, most (98%) of the error-suggesting variants were insertions or deletions (INDELs) or complex (i.e. a combination between single nucleotide polymorphisms (SNPs) and INDELs) (**Figure 2b**). In contrast, the Ref/Alt1 variants were mostly SNPs (**Sup. Fig. S1a**). Therefore, we proceeded to correct all error-suggesting variants, generating genome assembly Tb427v10. As expected, variant detection on the error-corrected Tb427v10 genome assembly showed a strong (98%) decrease in error-suggesting variants and a concomitant increase in Ref/Alt1 variants by ~5000, proportional to the number of Alt1/Alt2 variants corrected in Tb427v9. In total, we detected 96,060 heterozygote variants (**Figure 2b**), indicating an average of 4.21 heterozygote variants per kb in the core genome of *T. brucei* Lister 427.

**Figure 2.**
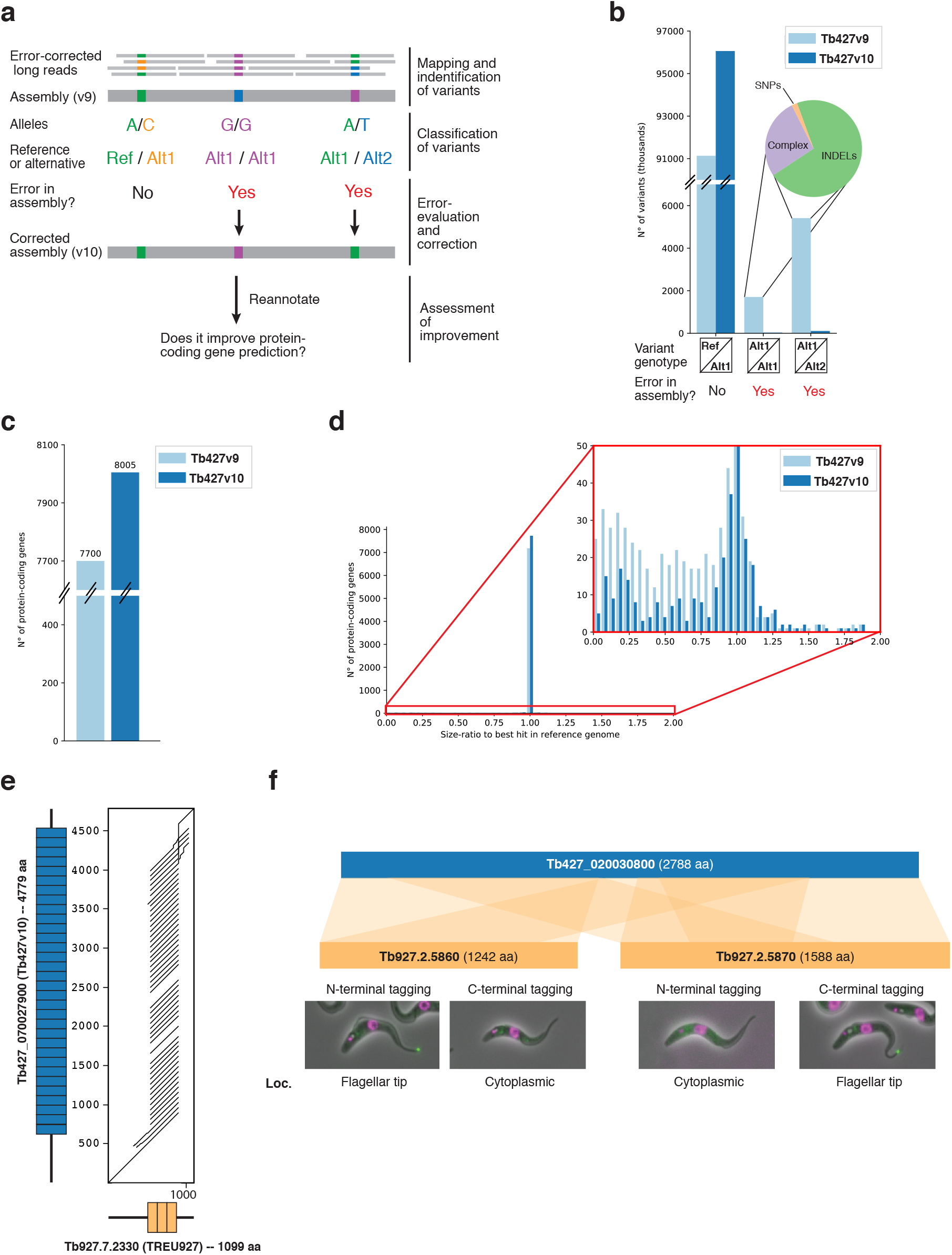
Genome error-correction. a) Error-correction approach. b) Number of variants identified before (Tb427v9) and after error-correction (Tb427v10) of the genome, grouped by variant genotype. The pie chart shows the proportion of SNPs, Complex and INDELs among the error-suggesting variants in Tb427v9. c) Number of protein-coding genes annotated. d) Size-ratio distribution of protein-coding genes compared to the *T. brucei* TREU927 genome assembly. The right panel is a zoom-in to the region selected in red. e) Alignment dot plot between synthenic orthologs in Tb427v10 (Tb427 070027900) and *T. brucei* TREU927 (Tb927.7.2330), showing a big repeat-array length difference, illustrated by the number of blue and orange boxes in the gene representations on the axes. f) Alignment of Tb427 020030800 from Tb427v10, to two contiguous protein-coding genes annotated in the *T. brucei* TREU927 genome assembly (Tb927.2.5860 and Tb927.2.5870), suggesting the gene was wrongly split in the latter assembly. The lower panel shows the subcellular localization (in green) of the N- and C-terminal tagged versions of Tb927.2.5860 and Tb927.2.5870 (data provided by TrypTag.org).

Next, to assess if the introduced sequence corrections were actually improving the genome assembly accuracy, we compared the annotation of protein-coding genes between the assembly versions. Given that most of the errors that we had corrected were INDELs or complex errors, we rationalized that if the error-suggesting variants were real errors (i.e. not technical artefacts), correcting them would ‘repair’ broken open reading frames, increasing the number of annotated protein-coding genes and decreasing the number of truncated protein-coding genes. The opposite effect would be expected should the corrections introduced errors.

Thus, we annotated the Tb427v10 assembly with the same pipeline used to annotate the Tb427v9 assembly. Comparing the two annotations, we observed an increase of 4% (305 genes) in the total number of annotated protein-coding genes in Tb427v10 (**Figure 2c**), indicating that the errors were real and that the correction step increased the accuracy of the genome assembly sequence. To further evaluate the quality of the genome and its annotation, we implemented a previously described strategy of ORF size comparison [22]. The strategy involves alignment of all annotated protein sequences from a query genome to a curated database of proteins followed by calculation of the size-ratio between each query protein and its best-hit in the database. If the query protein is of the expected length, the size ratio to its best-hit in the database will be ~1, if the protein in the assembly is truncated, the size ratio to its best-hit will be < 1. We compared the annotated proteome of the Lister 427 genome assembly, before (Tb427v9) and after (Tb427v10) the correction step to the proteome of the TREU 927 isolate of *T. brucei*, whose genome annotation has been curated extensively. Our comparison revealed a strong decrease in the number of genes with size-ratios below 1 (from 387 genes to 146 genes for size-ratio < 0.9) and an increase in the number of genes with size-ratio ~1 (from 7255 genes to 7761 genes for size ratio between 0.95 and 1.05) after the correction of the genome (**Figure 2d**), again supporting the importance of the error-correction approach.

Besides serving as a genome improvement assessment, the size-ratio analysis provided us with additional information. We identified genes where the size in our Lister 427 isolate assembly was larger than its best-hit in the TREU 927 isolate assembly (size-ratio >1). It is possible that these genes are larger in the Lister 427 isolate. However, a more plausible explanation seems to be that, given that in our assembly we used a long-read based approach, we could resolve the extension of repeat regions which were collapsed in the sanger-sequencing-based TREU 927 isolate assembly. Indeed, we analysed some of the cases and found examples of extreme repeat collapsing (**Figure 2e**) and incidents where our assembly suggests that two genes are in fact one (**Figure 2f**). The later finding is supported by microscopy data from the TrypTag project [23, 24]. N-terminal tagging of the first fraction of the gene (Tb927.2.5860) and C-terminal tagging of the second fraction of the gene (Tb927.2.5870) show a very specific localization in the flagellar tip of the parasite. While tags that break the actual gene (C-terminal tagging and N-terminal of Tb927.2.5860 and Tb927.2.5870, respectively) show cytoplasmic (mis)localization (**Figure 2f**, lower panel). The complete list of size-ratio between protein-coding genes from Tb427v10 and the TREU 927 isolate assembly is available as **Sup. Table S2**.

In summary, the usage of error-corrected PacBio reads led to an increase of 7% of protein-coding genes with the expected length, suggesting a significant improvement of overall genome sequence, and enabled the identification of ~96,000 heterozygote variants.

### Uneven distribution of variants across the genome

Following the improvement of the assembly accuracy, we proceeded to analyse the genome-wide distribution of the heterozygote variants identified. On average, we observed 4.21 variants per kb, yet variant distribution was very uneven, ranging from zero to more than 150 variants per 10 kb window (**Figure 3a-f** and **Sup. Fig. S4**). As expected, we obtained a very similar genomic distribution of variants from other sequencing datasets of different Lister 427-derived clones. Interestingly, we also observed a similar variant distribution in other *T. brucei* strains. Thus, we speculated the defined genomic distribution of variants to be caused by differences in mutation permissiveness, with non-coding regions, such as transcription start and transcription termination regions, possibly harbouring a higher variant density. However, we observed no clear correlation with any genomic feature analysed, such as H2A.Z and H3.V distributions, enriched at transcription start and termination sites, respectively [25] (**Sup. Fig. S2**).

**Figure 3.**
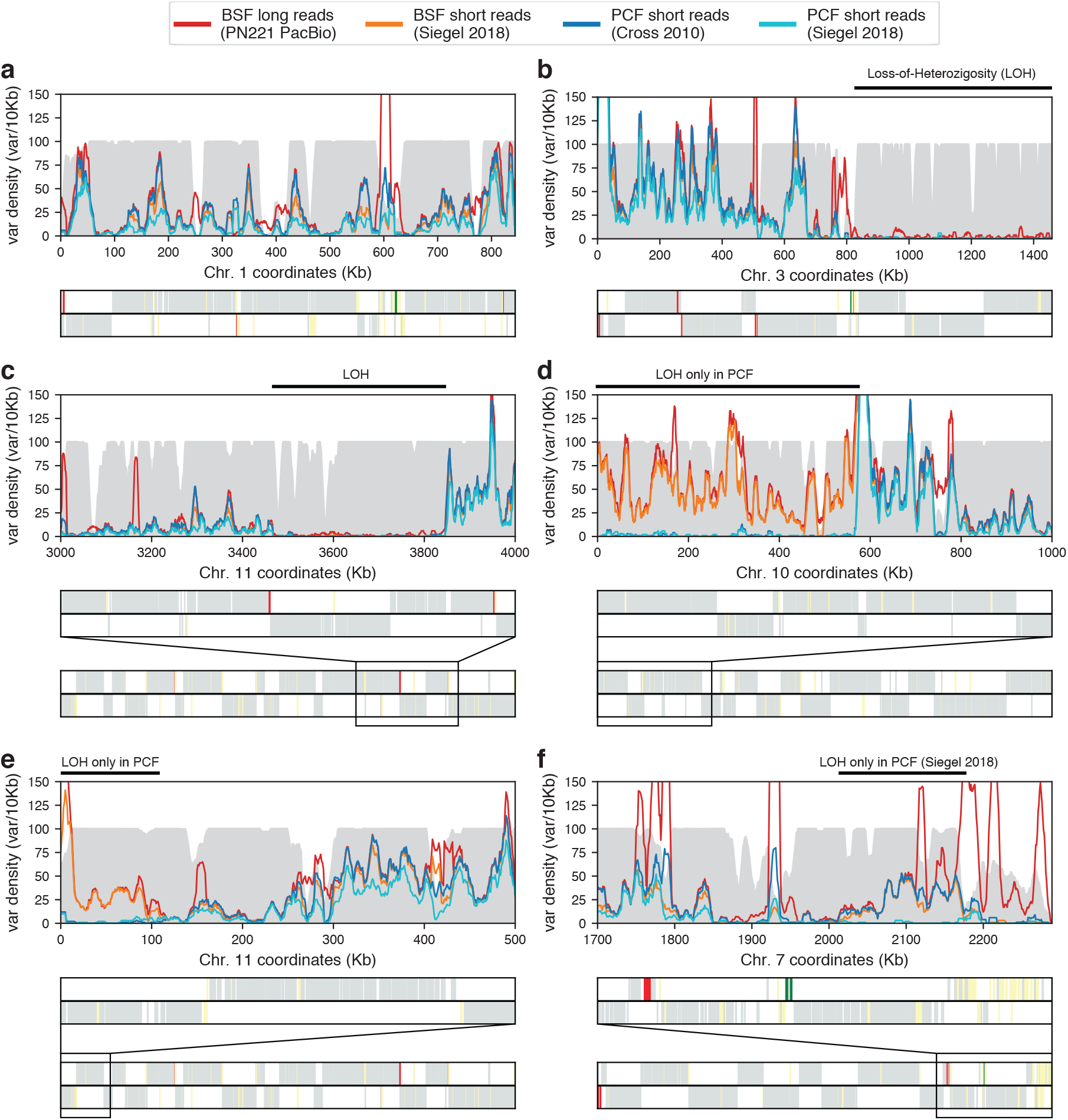
Variant density and loss-of-heterozigosity regions in the *T. brucei* Lister 427 genome. a-e) Heterozigosity from different *T. brucei* Lister 427 sequencing datasets on selected chromosomal regions of the Tb427v10 genome assembly. In the background, a filled-histrogram (gray) represents “mappability” (in range 0-100) for short-read sequencing data (see Methods). Loss-of-heterozigosity (LOH) regions are indicated by a black line above the plot. The lower panel shows the location and coding strand of protein-coding genes (grey), pseudogenes (yellow), VSG genes (red) and rRNA genes (green) as vertical lines, and Zoom-out representation of the complete chromosome when a smaller region was selected.

Noticeably, several chromosomes contained long regions where heterozygosity levels drop to near zero, representing loss-of-heterozygosity (LOH) regions. These regions were mostly located towards chromosome ends. For example, on chromosome 2 and 3 LOH regions extended for more than 300 and 600 kb inwards from the 3’ telomeres (**Sup. Fig. S4** and **Figure 3b**, respectively). On other chromosomes, LOH regions were also present internally (**Figure 3c**). To be able to exclude the possibility that LOH regions are the result of local haploidy, we analysed available DNA-seq data. No drop in coverage was observed for these regions that would have pointed to the partial loss of one of the two homologous chromosomes (**Sup. Fig. S3**). Thus, the observed LOH regions are most likely the result of recent recombination events between the two homologous alleles, which have reset heterozygosity.

To investigate how recent these recombination events may have occurred, we searched for LOH regions in different lines of the Lister 427 isolate. The *T. brucei* Lister 427 was isolated more than 50 years ago [26] and clones adapted to culture in mammal-stage condition or insect-stage condition have been selected early on [27]. As expected for cells that expand clonally in culture, we observed a very similar heterozygosity pattern across the chromosomes (**Figure 3** and **Sup. Fig. S4**). Most of the LOH regions were common to all clones tested. However, some were specific to insect-stage-adapted clones (**Figure 3d-e**) and one LOH region was only present in a specific insect-stage-adapted clone (**Figure 3f**). These observations suggest that most of the recombination events leading to LOH regions occurred before the cell-culture adaption, and that the insect-stage parasites are either more prone to recombination events or less sensitive to the, generally detrimental, loss-of-heterozygosity.

### Trisomy leads to proportional increase in transcription level

While analysing DNA-seq data to exclude haploidy in the LOH regions, we observed for a ‘wild type’ Lister 427 mammal-stage-adapted clone from our laboratory (“Tb427 BSF WT (Siegel 2014)”), originally acquired from the Cross laboratory, a ~50% increase in DNA-seq coverage across chromosome 5 compared to the remaining chromosomes (**Figure 4a-b**), pointing to a third copy of that chromosome. Other datasets derived from the same Lister 427 clone also showed increased coverage for chromosome 5, albeit some to a lower level, possibly the result of a mixed population. No increase in chromosome 5 coverage was observed for the cell-line used to assemble the Tb427v10 genome (“Tb427 BSF PN221”), that also derives from the Lister 427 isolate (**Figure 4a**), suggesting that the observed aneuploidy was incorporated during the in vitro culture after the generation of the Tb427 BSF PN221 cell line. To test if aneuploidy was common among *T. brucei* cell lines, we analysed available gDNA-seq datasets from other Lister 427 clones, from other *T. brucei brucei* isolates and from the related sub-species *T. brucei gambiense* and *T. brucei rhodesiense*. We did not observe any substantial changes in coverage between chromosomes for the datasets belonging to *T. brucei* TREU 927, *T. brucei gambiense* and *T. brucei rhodesiense* clones, suggesting normal ploidy in all the chromosomes (**Figure 4a**). Interestingly, we found two additional datasets from the *T. brucei* Lister 427 isolate suggesting aneuploidies, both from insect-stage-adapted clones. One showed increased coverage across chromosomes 2 and 6, while the other showed increased coverage across chromosomes 2 and 7 (**Figure 4a**), suggesting trisomy on these chromosomes. In all observed cases, the increase in coverage occurred across the entire chromosome (**Figure 4b-d** and **Sup. Fig. S5**), eliminating the possibility of partial chromosome amplifications. To our knowledge, aneuploidy has not been previously reported in *T. brucei*.

**Figure 4.**
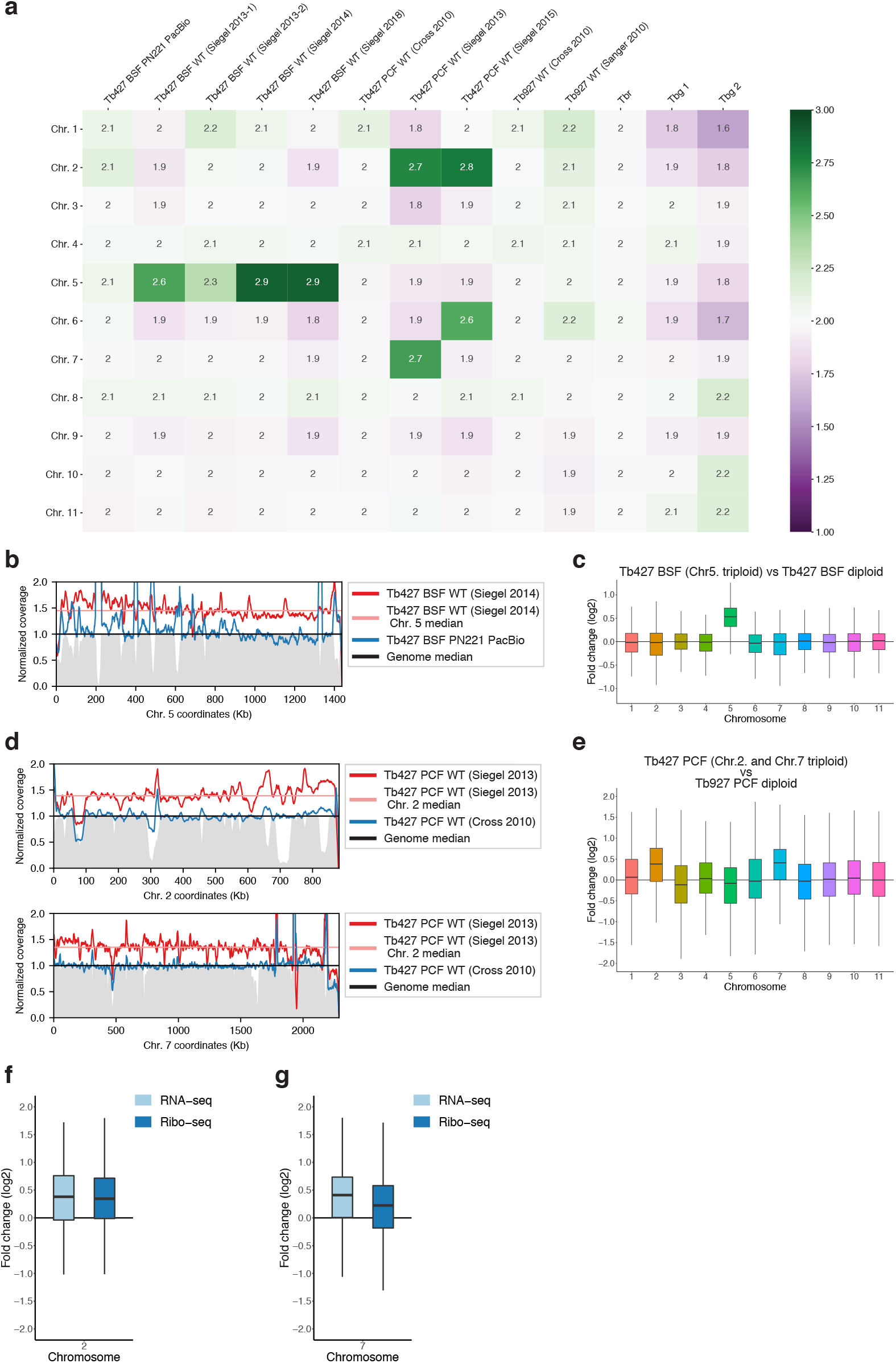
The effect of aneuploidy in gene expression in *T. brucei*. a) Median coverage per chromosome for different *T. brucei* DNA-seq datasets, normalized to genome median and centered at two (to illustrate expected diploidy). In the labels, “Tb427” indicates a Lister 427 derived clone, “Tb927” a TREU927 derived clone, “Tbr” *T. brucei rhodesiense* and “Tbg” *T. brucei gambiense*. b) Normalized coverage density in chr 5 for Tb427 BSF WT (Siegel 2014) clone (dark red line) and its median (straight light red line) compared to Tb427 BSF PN221 PacBio clone (dark blue line). The genome median is set to 1 (straight black line). Mappability is shown as a grey filled-line (in range 0-1). c) RNA-seq fold change (log2) pooled by chromosome, from a *T. brucei* Lister 427 clone triploid for chr 5 over a diploid *T. brucei* Lister 427 clone. d) Normalized coverage density in chr 2 (upper panel) and Chr. 7 (lower panel) for Tb427 PCF WT (Siegel 2013) clone (dark red line) and its median (straight light red line) compared to Tb427 PCF WT (Cross 2010) clone (dark blue line). The genome median is set to 1 (straight black line. Mappability (in range 0-1) is indicated in grey. e) RNA-seq fold change (log2) pooled by chromosome, from a *T. brucei* Lister 427 PCF triploid for chr 2 and chr 7 over a diploid *T. brucei* TREU927 clone. f) and g) RNA-seq and Ribo-seq fold change (log2) between the same clones as e), for chr 2 and chr 7, respectively.

The organization of genes in polycistronic transcription units [16] and the lack of canonical RNA pol II promoters [28] has led to the assumption that RNA pol II transcription is not regulated in *T. brucei*. If true, the increase in chromosome copy number should thus result in a proportional increase in transcript levels. To test this assumption, we compared available RNA-seq data from a Lister 427 clone with trisomy 5 to a diploid clone carrying two copies of chromosome 5. Our analysis showed a ~50% increase in the transcript level for the genes in chromosome 5 in the cell line with trisomy 5 (**Figure 4c**), indicating a direct correlation between gene copy number and transcriptional level. These observations support the assumption that *T. brucei* lacks transcriptional regulation, but also suggests the absence of a general dosage compensation mechanism that senses and regulates transcriptlevels post-transcriptionally.

Similar results were obtained when comparing RNA-seq data from the insect-stage adapted Lister 427 clone [29] containing trisomy 2 and 7 to RNA-seq data from a regular diploid *T. brucei* TREU 927 isolate insect-stage adapted clone [30] (**Figure 4e**). Interestingly, comparing ribosome profiling data from both clones, we observed a negligible reversion of expression levels (textbf Figure 4f-g), suggesting minimal feed-back regulation at the translational level.

### Haplotype-phasing of the Lister 427 genome assembly

With the knowledge of the uneven genome-wide variant distribution and having confirmed diploidy for all chromosomes in the clone used for the genome assembly, we went ahead to generate an allele-specific genome assembly of *T. brucei* Lister 427. To this end, we combined raw PacBio reads and Hi-C data, together with HapCUT2 [15], a tool capable of integrating information from diverse sequencing technologies to link variants into haplotypes. Following this approach, we could assign 99.55% of the total heterozygote variants (95628/96060 variants) to a specific chromosomal haplotype, enabling the almost complete reconstruction of the two alleles for each chromosome ‘core’ (**Figure 5**).

**Figure 5.**
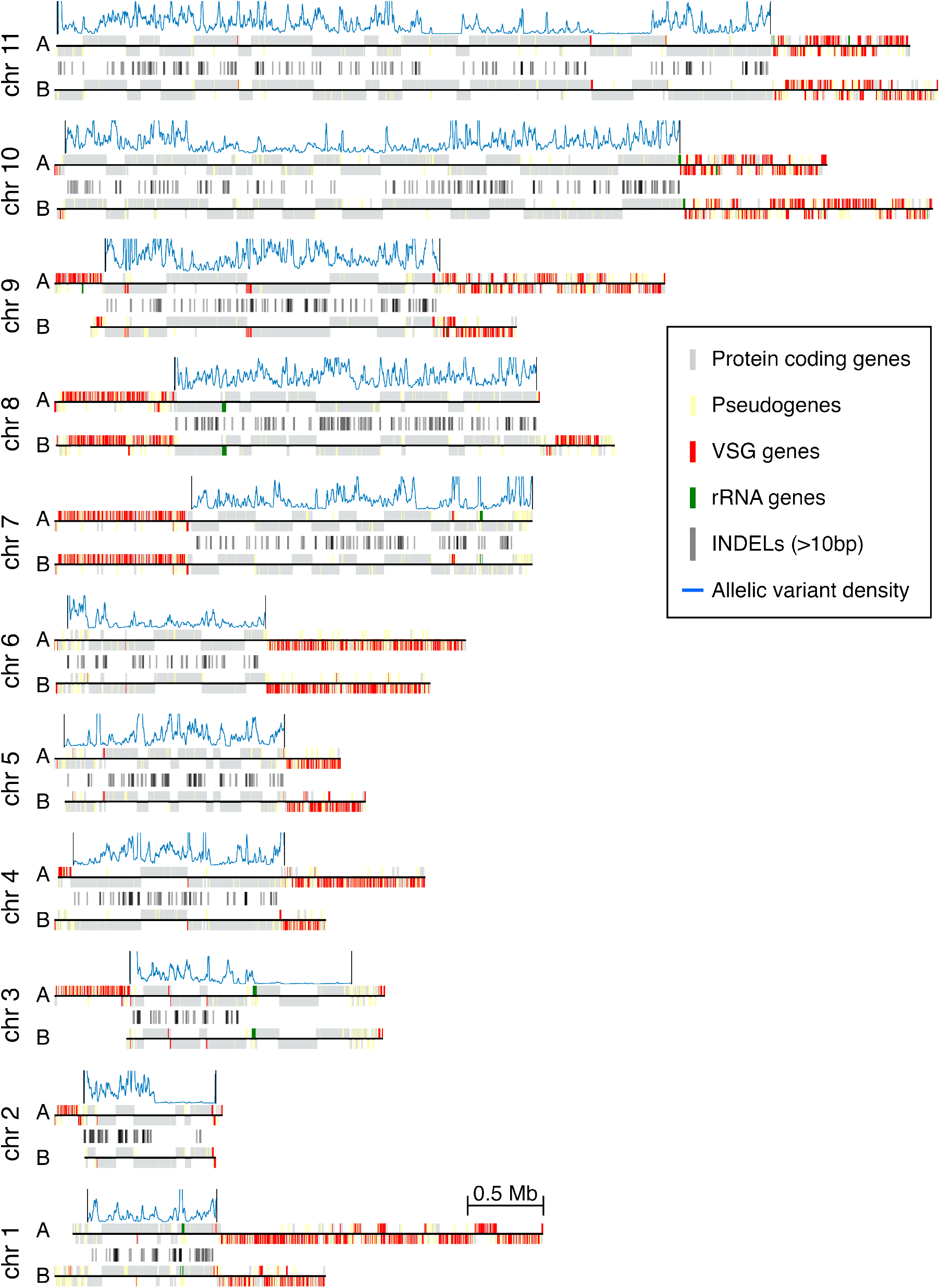
Fully phased *T. brucei* Lister 427 genome. Both alleles are plotted for each chromosome. Protein-coding genes (gray), pseudogenes (yellow), VSG genes (red) and rRNA genes (green) are indicated by vertical lines on top or bottom (depending on the coding strand) of the black lines representing the chromosomal sequence. Variant density is indicated with a blue histogram on top of each chromosomal ‘core’ region. INDELs (>10bp) are indicated by dark grey lines between the ‘cores’.

To evaluate the accuracy of the haplotyping and to reconstruct the full chromosome alleles, we quantified the Hi-C interactions between the chromosome ‘core’ alleles and the haploid-like subtelomeres. We observed that each alternative subtelomere interacted strongly with only one core allele, in a distant-dependent fashion (**Sup. Fig. S7**). This allowed us to scaffold the full chromosome alleles (**Figure 5**), and also served as validation of the phasing approach. An inaccurate variant phasing of the chromosomal cores would have led to indistinguishable or patched interactions between the haploid-like subtelomeres and both core alleles, which was not observed.

### Allele-specific premature termination codons affect translation but not transcription level

To investigate the biological significance of haplotype variations, we annotated each of the haplotypes independently and characterized a selection of allelic differences. In total, we detected 8030 and 8007 genes for haplotypes A and B, respectively. Both haplotypes contained more annotated genes than the collapsed Tb427v10 genome assembly (**Sup.Fig. S1b**, already suggesting that haplotype phasing fixed errors present in the collapsed assembly. In order to identify specific differences in the annotated proteome between haplotypes, and in comparison to the collapsed Tb427v10 assembly, we performed reciprocal all vs all BLAST between the three assemblies and analysed the size-ratio to best-hits, similar to our analysis following the initial error-correction.

First, we aimed to understand why some gene ORFs were of the expected length only after haplotyping and took a closer look at some exemplary genes. We found that one reason for ORF truncations in the collapsed assembly was the presence of internal repeats of varying length between the alleles, leading to frame-shifts during collapsing. Another apparent reason for ORF truncation in some genes was the presence of nearby allelic variants which, by chance, generated a shifted sequence homology between the alleles, leading to wrong read alignments and the introduction of INDEL errors in the collapsed assembly (For an example, see **Sup. Fig. S8**). These findings highlight the importance of allele phasing for the accurate identification of protein-coding genes in heterozygous organisms.

We next wondered if there were allelic variants introducing premature termination codons (PTCs) in protein-coding genes, leading to different CDS length between the alleles. We analysed the reciprocal BLAST results searching for genes for which the CDS in one of the alleles was more than 10% smaller or was directly absent. Based on this analysis we identified 123 putative genes with different CDS length between the alleles (**Sup. Table S3**).

Finally, to investigate the implications of PTCs on gene expression, we analysed allele-specific transcription and translation for genes with an allele-specific PTC. We used RNA-seq and ribosome profiling data and quantified the number of reads mapping to each allele for each variant position in the genes. Two examples are illustrated in **Figure 6**. Gene Tb427 090051600 (v10), annotated as hypothetical protein, encodes a protein of 379 aa in haplotype “B”, identical to its orthologs in other *T. brucei* strains, while haplotype “A” contains a 1 bp insertion shortly after the start of the CDS, which changes the reading frame and leads to a short ORF (71 aa) followed by a second ORF (300 aa) just downstream. The second ORF contains a complete functional domain, based on InterPro[31] potentially coding a “P-loop containing nucleoside triphosphate hydrolase”. Our allele-specific analysis expression analysis indicates that both alleles are fully transcribed, but that for haplotype “A” only transcripts from the first ORF are being translated, leading to a short, probably non-functional, polypeptide (**Figure 6a**).

**Figure 6.**
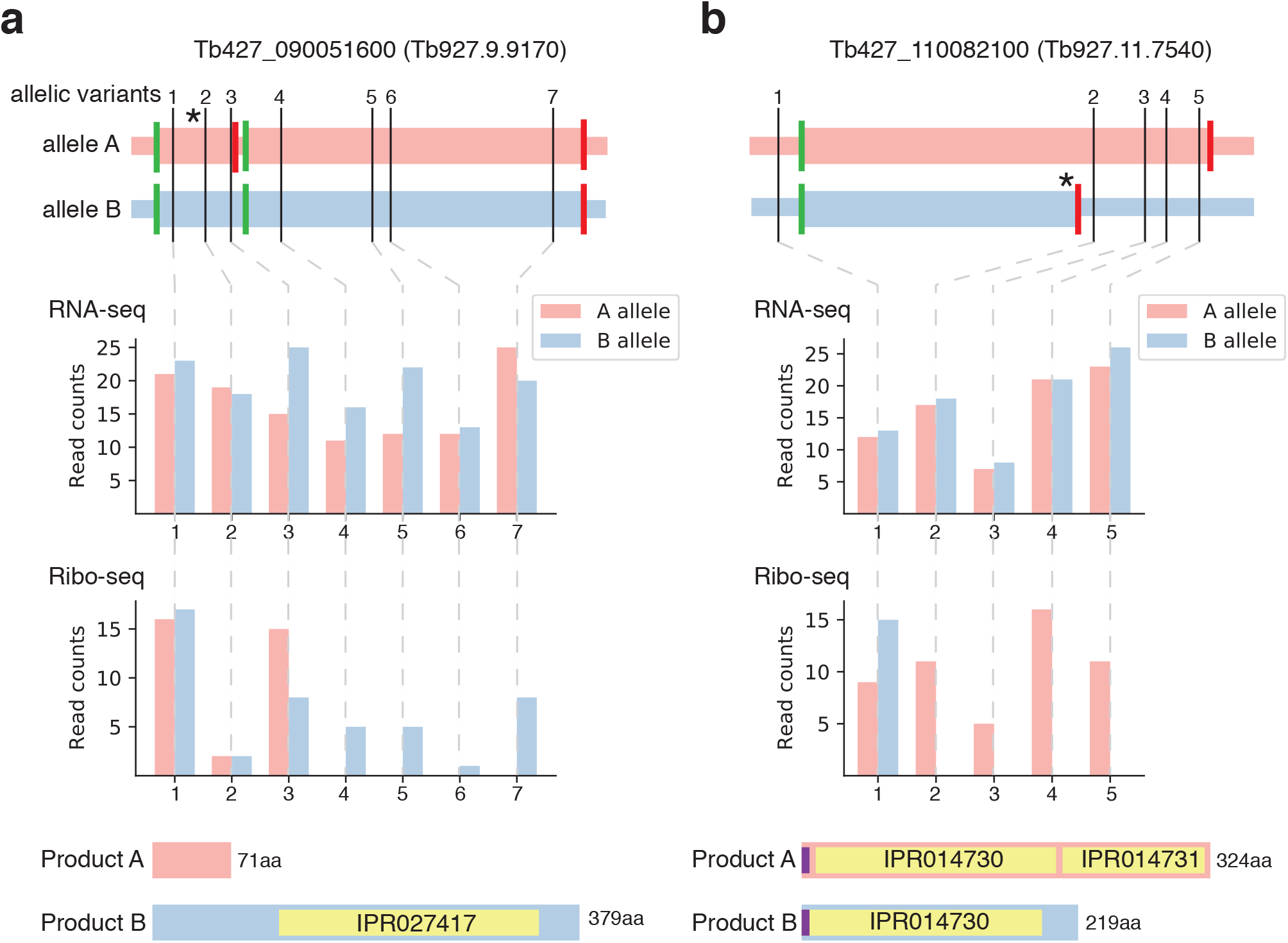
Allele-specific transcription and translation in genes with allele-specific premature termination codons. a) and b) Examples of genes with allele-specific variants leading to a premature termination codon. In the upper panels the thiner pink and blue horizontal lines represent the sequence of the two alleles, while the thicker line on top indicates the open reading frames. Variant positions are indicated by black vertical lines and numbered. An asterisk indicates the position of a frameshifting INDEL variant. In green and red vertical lines, start and stop codons are indicated, respectively. The middle panels show RNA-seq and Ribo-seq read counts for both alleles in each of the variant positions. The lower panels show the expected protein size for each allele. Yellow boxes indicate InterPro domains and violet boxes signal peptides.

Another example is the gene Tb427 110082100. It encodes a putative electron-transfer flavoprotein alpha of 324 aa in haplotype “A”, identical to its orthologs in other *T. brucei* strains. On haplotype “B” the gene has a 1bp insertion towards the middle of the actual ORF, leading to a shorter ORF of 219 aa, without any other possible start codon downstream. The quantification of the reads on the variant positions indicates that both alleles are being transcribed at similar levels. However, as expected, translation of the 3’ region only occurs for transcripts of haplotpye “A” (**Figure 6b**). It seems likely, that haplotype “B” is producing a truncated protein containing the signal-peptide and the first flavoprotein domain (IPR014730). Interestingly, electron-transfer flavoproteins are heterodimers composed of an alpha and a beta subunit [32]. The alpha subunit contains the domain IPR014730 and IPR014731, while the beta subunit contains only the IPR014730 domain. Thus, our observations raise the possibility that the truncated protein encoded by allele B, mimics the beta subunit of the electron-transfer flavoprotein.

In summary, for these two genes, as well as for additional 4 genes with PTCs (**Sup. Fig. S9**), we observed that transcript levels are unaffected by PTCs, while ribosome occupancy drops directly downstream PTCs and there is no indication of translation being reinitiated.

### Location and completeness of the VSG gene repertoire

*T. brucei* harbours a vast repertoire of antigenic genes, named Variant Surface Glycoproteins (VSGs) [33]. While most of the VSG repertoire is located in subtelomeric arrays in the eleven chromosomes [17], part of it is located in so-called minichromosomes, 30-150 kb long chromosomes [34]. What roles the VSG repertoires in different contexts, and specifically in minichromosomes, play in antigenic variation remains poorly understood. To facilitate these types of studies, we decided to identify minichromosomes among the unassembled contigs. We searched for specific minichromosomal signatures [35] and for minichromosomal VSGs previously identified in this strain [36], leading to the identification of 60 minichromosomal contigs. On them we found a total of 51 VSGs, all of them located in different contigs, except for one case where we found two VSGs on the same contig. The number of VSG genes in the different genomic locations of Tb427v10 genome assembly is shown in (**Table 1**. Previous studies based on different genome assemblies indicate that a very low portion of the VSG repertoire is fully functional, i.e. most of them are pseudogenes or gene fragments [16, 37, 36]. We analysed the 2872 VSG genes annotated, searching for the presence of required features for a VSG to be functional (e.g. peptide signal, GPI-anchor signal), and found that only 433 (15%) can be considered as fully functional, in line with previous estimations for *T. brucei* Lister 427 [36]. Interestingly, as shown before, we found that minichromosomes are enriched in fully functional VSGs (43 out of 51, 84%). The complete list of VSG genes, their location in the genome and which ones are predicted fully functional is available in **Sup. Table S4**.

**Table 1:** VSG location and completeness. Number of VSGs in the different genomic locations in the *T. brucei* Lister 427 genome assembly, and the number of VSGs predicted to be fully functional.

| Location | N° of VSGs | Fully functional |
| --- | --- | --- |
| Subtelomeric | 2634 | 346 |
| Chromosomal core | 27 | 3 |
| Minichromosomal | 51 | 43 |
| BES (Telomeric) | 13 | 13 |
| BES (Non-telomeric) | 4 | 0 |
| MES | 8 | 7 |
| Unassigned contig | 135 | 21 |
| Total | 2872 | 433 |

## Discussion

The most frequent errors in long-read sequencing technologies are insertions and deletions [19]. This has been shown to particularly affect the accuracy of protein-coding gene annotation, by introducing frameshifts and premature stop codons [20]. Here, we implemented an error-correction step to the *T. brucei* Lister 427 genome assembly using short-read error-corrected PacBio reads. To evaluate the relevance of the correction process, we compared protein-coding gene annotation before and after correction. We observed a considerable improvement, correcting more than 500 protein-coding genes, representing 7% of the whole proteome. Our results suggest that the application of error-correction steps is crucial to achieve high-quality genome assemblies and that, as has been shown before [38], measuring protein annotation accuracy is a very informative way to assess it. We believe that a process similar to the one we performed should be applied to correct available non-hybrid genome assemblies from long read technologies. In the meantime, caution should be taken when analysing protein-coding genes in long-read based assemblies, especially when observing apparently absent genes, pseudogenes, or truncated genes compared to closely related organisms. For future genome assemblies, this issue could be considered from the beginning by either implementing a hybrid strategy in the assembly (e.g, generating short-read error-corrected long reads and use them as input in the assembly) or using the latest HiFi reads from PacBio, which have shown to reduce INDEL errors [39]. The average heterozygosity of the *T. brucei* Lister 427 was 4.2 variants / kb. This is almost twice the number predicted for *T. brucei* TREU 927 (2.2 variants / kb) [40]. In comparison, heterozigosity in *Leishmania* is in general very low [41], ranging from almost complete homozygosity to 1.6 variants / kb in an extremely heterozygous outlier from *Leishmania donovani* [42]. In *Trypanosoma cruzi* heterozygosity was shown to be around 2 variants / kb [43], with the exception of strains belonging to hybrid lineages, where heterozygosity can reach levels of 20 variants / kb [44].

Importantly, the average heterozygote variant density within CDSs in the *T. brucei* Lister 427 strain was 2.5 variants / kb, and almost 50% of the CDSs had two or more phased variants (>25% had four or more), making our phased genome a very interesting template to study genome-wide allele expression bias in kinetoplasts. A phased genome with similar variant density [45] was successfully used to study allele-specific expression control in the fungal pathogen *Candida albicans* [46].

Long regions with LOH were identified in different chromosomes of the Lister 427 genome. Most of them were conserved across clones adapted to mammal-stage and insect-stage, suggesting that the recombination events leading to them occur prior to clonal adaptation. LOH regions are thought to arise after “gene conversion” compensatory mechanism for counteracting deleterious mutations in asexual species [47]. However, all sub-species of *T. brucei*, with the exception of the human infective *T. brucei gambiense* Group 1, have the ability to undergo sexual reproduction [48]. Thus, the high number of LOH regions identified in the Lister 427 genome is unexpected. A plausible explanation is that the recombination events leading to the LOH regions occurred in the early phases of cell culture, possibly playing a role in the adaptation to laboratory conditions. An ancestral line of present Lister 427 clones was shown to be fly transmissible and able to undergo meiosis and genetic exchange [26, 48], while all the Lister 427 derived clones we analysed are monomorphic, i.e. lost the ability to differentiate into other forms. Sequence comparison among this ancestral line and the current ones might shed light on the origin of the LOH regions and allow to pinpoint the genetic changes responsible for losing the ability to complete the natural development cycle.

We identified *T. brucei* Lister 427 clones with trisomy in one or two of the eleven chromosomes. This is, to our knowledge, the first time that aneuploid *T. brucei* clones have been described. Chromosomal aneuploidy was shown to be extensive in other Kinetoplastids, such as *Leishmania* [49, 42, 50] or *T. cruzi* [51], but there was still no evidence that this could occur in Kinetoplastida from the Salivarian evolutionary branch. However, given that we only analysed clones adapted to laboratory conditions, we cannot rule out that the aneuploidies observed are a laboratory artefact, not occurring in the natural life cycle of *T. brucei*. Indeed, recent efforts analysing gDNA-seq data from multiple field isolates of *T. brucei* could not detect any aneuploidy [52]. Despite the fact that the observed aneuploidies could be only an adaptation to laboratory conditions, it is interesting that different chromosomal trisomies were selected in clones adapted to different life cycle stages. Stage-specific aneuploidies were previously observed in *L. donovani* [53]. This suggest that either genes located in distinct chromosomes promote growth in the alternative life cycle stages or that the tolerance for over-expression of genes in distinct chromosomes is different or, probably, a combination of both. Data from other culture-adapted clones and field isolates may shed light on this matter.

The availability of transcriptomic data from aneuploid clones allowed us to assess the impact of ploidy on transcription. We observed a proportional increase of mRNA transcript level throughout the trisomic chromosomes, supporting the absence of transcriptional regulation, a general dosage compensation mechanism and global allele-specific expression in *T. brucei*. It would be interesting to analyse quantitative proteomic data in the aneuploid clones to see if there is a compensatory mechanism at this stage or if the parasite can deal with chromosome-wide 50% increase in protein level. In *Leishmania* parasites, several studies have also shown an overall correlation of ploidy and transcription [53, 54, 55], albeit with gene dosage-independent fluctuations for some chromosomes and indications of regulatory mechanisms compensating at the protein expression level[55]. On a practical matter, the observation that there is a direct correlation of mRNA transcript level and ploidy, indicates that comparative RNA-seq data could be used to directly determine deviations from euploidy in *T. brucei* field isolates.

There was an increase in protein-coding genes annotated for each haplotype after phasing in comparison to the collapsed assembly (Tb427v10), suggesting that this procedure is also important for accurate identification of protein-coding genes. Indeed, we identified several single copy genes that were only corrected after phasing. Furthermore, we identified gene alleles encoding protein of different sizes and using the haplotype information in combination with RNA-seq and ribosome profiling data we could validate the predicted size-differences at the translational level, illustrating the usefulness of generating haplotype specific assemblies to study allele-specific behaviours.

In many eukaryotes, mRNAs containing premature termination codons (PTCs) are rapidly degraded by a process named “nonsense-mediated decay”(NMD). Authors of a previous study suggested that this mechanism was also active in *T. brucei* [56], given that they observed a reduction of mRNA after introducing PTCs in an endogenous gene. Here, in all the genes we analysed with one allele containing a PTC, we see no evidence of NMD, i.e. transcript levels remain even for both alleles, even for genes where the PTC occurs very early on, arguing against the presence of fully active NMD in *T. brucei*.

We believe that the availability of an accurate allele-specific genome assembly of *T. brucei* will serve as a better platform for the design of constructs in gene-specific studies but will also allow to address multiple questions at the genome-wide level. Especially in combination with single-cell RNA-seq approaches, it will be possible to address questions about transcriptional kinetics or allele-specific expression. Furthermore, the first allele-specific genome assembly of *T. brucei* will serve as a benchmark in the development of much needed fully-automatic pipelines for the genome assembly of field isolates with extreme levels of heterozygosity in order to fully understand the diversity and evolution of this parasite. The development of such pipelines will not only be useful for the assembly of trypanosomes genomes, but in general for animal or plant wild isolates with high-levels of heterozygosity.

## Methods

### Error-correction of PacBio reads

Error-corrected PacBio reads were generated with Proovread [21] using as input PacBio raw reads and a set gDNA-seq short-reads from *T. brucei* Lister 427 (see **Sup. Table S1**).

### Variant detection

Error-corrected PacBio reads were mapped to the genome assemblies with Minimap (v2.10) [57], while short-reads were mapped with bwa-mem (v0.7.16) [58]. Supplementary, secondary and low-quality alignments (mapq < 10) were filtered with samtools (v1.8) [59]. Variant detection was performed with FreeBayes (v1.2.0) [13], and filtered by QUAL > 20.

### Assembly error-correction

Variants suggesting errors (i.e. homozygous for an alternative allele, or heterozygous for two alternative alleles) in the *T. brucei* Lister 427 v9 genome assembly identified from mapping error-corrected PacBio reads, were corrected using a Perl (v5.26.2) script parsing the vcf file. This error-correction step lead to Tb427v10 genome assembly.

### Annotation

Genome assembly versions were annotated primarily with Companion v1.0.1 [60]. Additionally, Variant Surface Glycoprotein (VSG) genes were annotated based on BLAST results to the manually curated VSGnome [36], as done previously [17]. Genes with annotated introns were considered pseudogenes, and the annotation modified.

### Comparison of annotated proteome

Coding sequences from the annotated protein-coding genes from the chromosomal cores for each genome assembly version were extracted with BEDTools (v2.26.0) [61] ‘getfasta’ function, and translated with fastaq (v3.17.0) ‘translate’ function. Each proteome multifasta was reciprocally aligned and to the proteome from *T. brucei* TREU927, (version 42, extracted from TriTrypDB [62]) using BLAST (v2.7.1+) [63]with settings to generate a tabular extended output “-outfmt ‘7 std qlen slen’”. For each query protein, the best-hit in the subject proteome was extracted based on the maximum Bitscore. The analysis of query-size over best-hit protein-size was done using python (v3.6), with the module pandas (v1.03) [64] and the plots were done using matplotlib (v3.2.1) [65].

### Density distribution analysis

The *T. brucei* Lister 427 genome assembly v10 (Tb427v10) was divided in 10 kb overlapping bins, with 1 kb sliding window with BEDTools (v2.26.0) ‘makewindows’ function. For each DNA-seq dataset, the number of heterozygote variants within each bin was counted with BEDTools ‘coverage’ function. For the same bins, the number of mapped reads for the different sequencing datasets analysed (See **Sup. Table S1**) were counted. GC-content was calculated with BEDTools “nuc” function. ‘Mappability’ was defined as the percentage of mapping reads obtained with reads filtered by mapping quality (mapq > 10) from the mapping reads obtained without filtering. Mappability serves as an indicator on how repetitive (low mappability) or unique (high mappability) region are. Pearson correlation between the different datasets was calculated with the module pandas from python and the heatmap was generated with the module seaborn (v0.10.0) [66].

### Ploidy analysis

To estimate the ploidy for each chromosome in each gDNA-seq datasest, the median coverage was divided by the median coverage for all the chromosomes. For the specific analysis of candidate chromosomes with trisomy, their median coverage was divided by the median coverage of the rest of the chromosomes.

### Differential expression analysis

RNA-seq and ribosome profiling data were mapped to Tb427v10 genome assembly with bwa-mem (v0.7.16), converted to bam files with samtools (v1.8). Gene counts were obtained with subread (v1.6.2) [67] function ‘featureCounts’ and differential expression analysis was performed in R (v3.5.2) with DESeq2 (v1.22.2) [68]. Box plots of log2-fold-change for each chromosome were generated using ggplot2(v3.3.0).

### Allele phasing

The heterozygote variants identified in the diploid ‘core’ of Tb427v10 genome assembly were phased with HapCUT2 [15] using raw PacBio reads and Hi-C data as input with settings to phase not only single-nucleotide variants but also INDELs (“–indels 1”). The phasing output of HapCUT2 was used to generate the allele “A” and the allele “B” fasta genome file using a custom-made perl script. HapCUT2 generates phased “blocks” for each chromosome. In case of several blocks, only the largest block was considered.

### Minichromosomal VSG identification

To identify putative minichromosomal contigs, we first searched for contigs containing a curated set of minichromosomal VSGs [36]. At the same time, we search for contigs containing a minichromosomal signature, a 177 bp repeat [35]. We selected two regions of the 177 bp repeat which were highly conserved [69], performed BLAST searches and extracted the contigs which had at least one hit. The search with both regions returned us the same set of contigs. Additionaly, we searched for contigs containing telomeric repeat (we used five times ‘TTAGGG’ as query in the BLAST search) and a VSG annotated. Finally, we combined the set of putative minichromosomal contigs identified with the different approaches.

To assess the completeness of VSGs, we first generated a protein multifasta of all annotated VSGs combining BEDTools ‘getfasta’ function and fastaq ‘translate” function. We predicted the presence of peptide signal using SignalP (v5.0) [70] and GPI-anchor using netgpi (v1.1) [71] and determined loose VSG protein size limits (between 400-600 aa) based on size ranges previously observed [36]. Given that we noticed that frequently start and stop codons were not correctly annotated for the VSGs (e.g. CDS annotation was starting after the ‘ATG’ and finishing before the stop codon), we decided to make a more sensitive selection of CDSs. We extracted from each annotated VSG its coordinates extending 300 bp on each side. Then, we determined the longest ORF and perform the same predictions as before. A VSG was determined as functionally complete if it had (either in the standard or in the sensitive approach) a predicted peptide signal, a predicted GPI-anchor and a size between 400-600 aa.

### Hi-C interactions to the fully-phased genome

Hi-C reads were truncated with HiCUP (v0.8.0) [72], mapped independently with bwa-mem to the *T. brucei* Lister 427 allele-specific genome assembly. Normalized interaction frequency matrices were generated using HiC-Pro (v2.11.4) [73] and heatmaps were obtained with HICsuntdracones (v0.2.0), following the same pipeline we have used previously [17].

### Allele-specific quantification

Allele-specific read counts for RNA-seq and ribosome profiling data from *T. brucei* Lister 427 [30] were obtained using GATK (v4.1.8.1) [14] function ‘ASEReadCounter’ with ‘–min-mapping-quality 10’, and the variant file generated for the error-corrected PacBio reads on Tb427v10 (obtained in ‘variant detection’ section), filtered to contain only SNPs. Then, the counts for each allele were assigned to haplotypes based on the HapCUT2 phasing output, using a python pandas script. Bar plots for the allelic counts in selected genes with premature termination codons were done using matplotlib.

### Published dataset used

Published datasets used in this study are described in **Sup. Table S1**, available in the Zenodo repository dedicated to this article (https://doi.org/10.5281/zenodo.4674608).

## Data availability

All the new sequencing data is available in the European Nucleotide Archive (ENA) under the primary accession number PRJEB43606. Whereas the Tb427v10, the Tb427v10 phaseA and the Tb427v10 phaseB, the Tb427v10 phased diploid, and their respective annotation files can be found in Zenodo (https://doi.org/10.5281/zenodo.4674608). The Tb427v10 genome assembly is already integrated into TriTrypDB [62], the new datasets will be made available soon. All the supplementary tables are also found in the Zenodo repository.

## Code availability

Workflows and custom-made Unix Shell, Perl, Python and R scripts have been deposited to Zenodo (https://doi.org/10.5281/zenodo.4674608).

## Acknowledgments

Localization data for Tb927.2.5860 and Tb927.2.5870 were provided by the TrypTag project (TrypTag.org). We would like to thank Frank Förster (University of Würzburg, Germany) for generating the error-corrected PacBio reads. We also would like to thank the Bioinformatics Core Facility from the Biomedical Center at LMU Munich for access to the computing cluster, Tobias Straub and all members of the Siegel laboratory for valuable discussions, Fernán Agüero, Vanessa Luzak and Kirsty McWilliam for critical reading of the manuscript. This work was funded by an ERC Starting Grant (3D Tryps 715466) to T.N.S. R.O.C was supported by a Georg Forster Fellowship (Humboldt Foundation).

---

**Figure S1.**
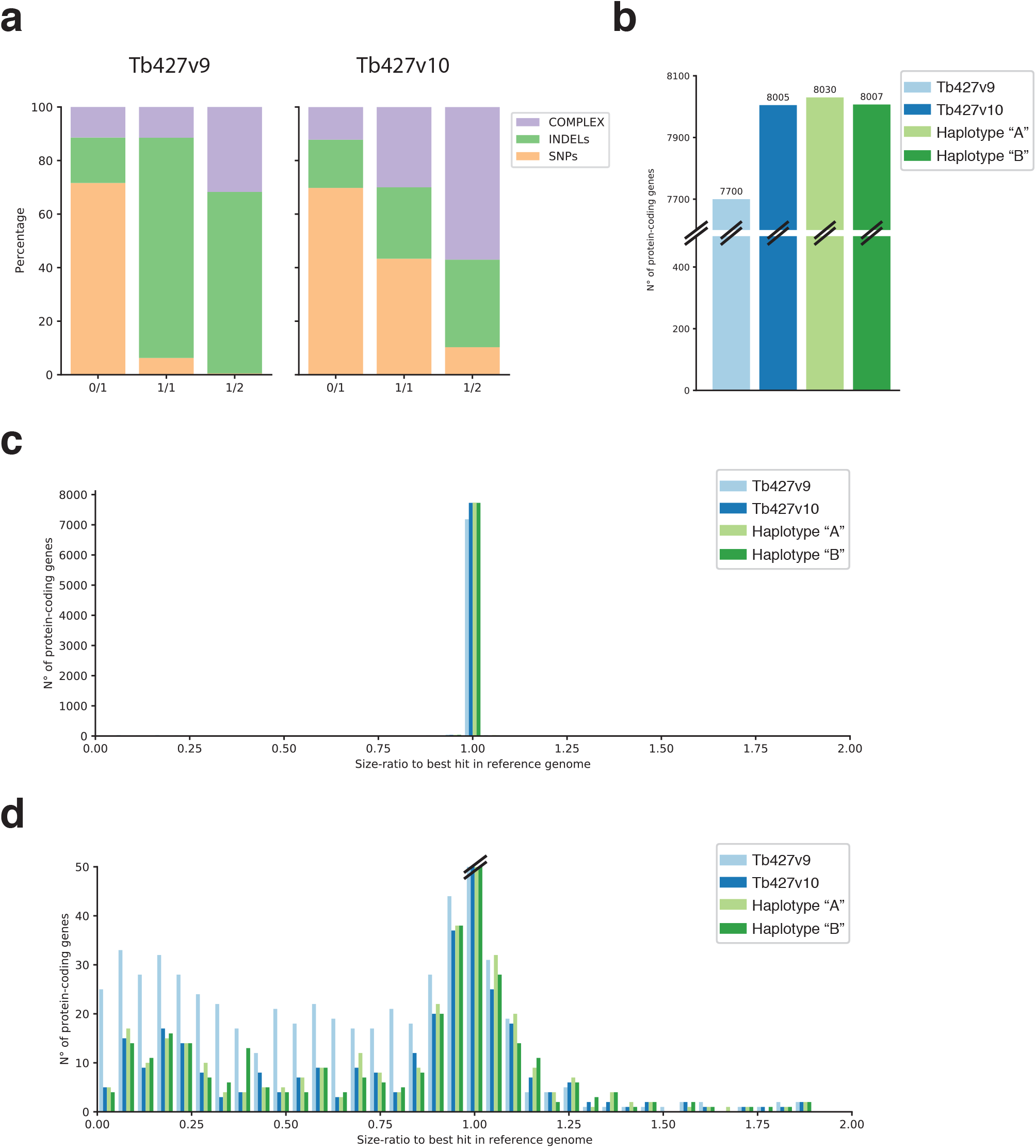
Variant proportion and assessment of protein content. a) Proportion of SNPs, INDELs and Complex (i.e. combination between SNPs and INDELs) variants for the different genotype scenarios (Ref/Alt1 (0/1), Alt1/Alt1 (1/1), Alt1/Alt2 (1/2)). B) Protein-coding genes annotated for the different *T. brucei* Lister 427 genome assemblies (Tb427v9, Tb427v10, haplotype “A” and haplotype “B”). c) Histogram of size-ratio of protein-coding genes to the best hit in *T. brucei* TREU927 genome for the different *T. brucei* Lister 427 genome assemblies. D) Zoom-in to the plot in c).

**Figure S2.**
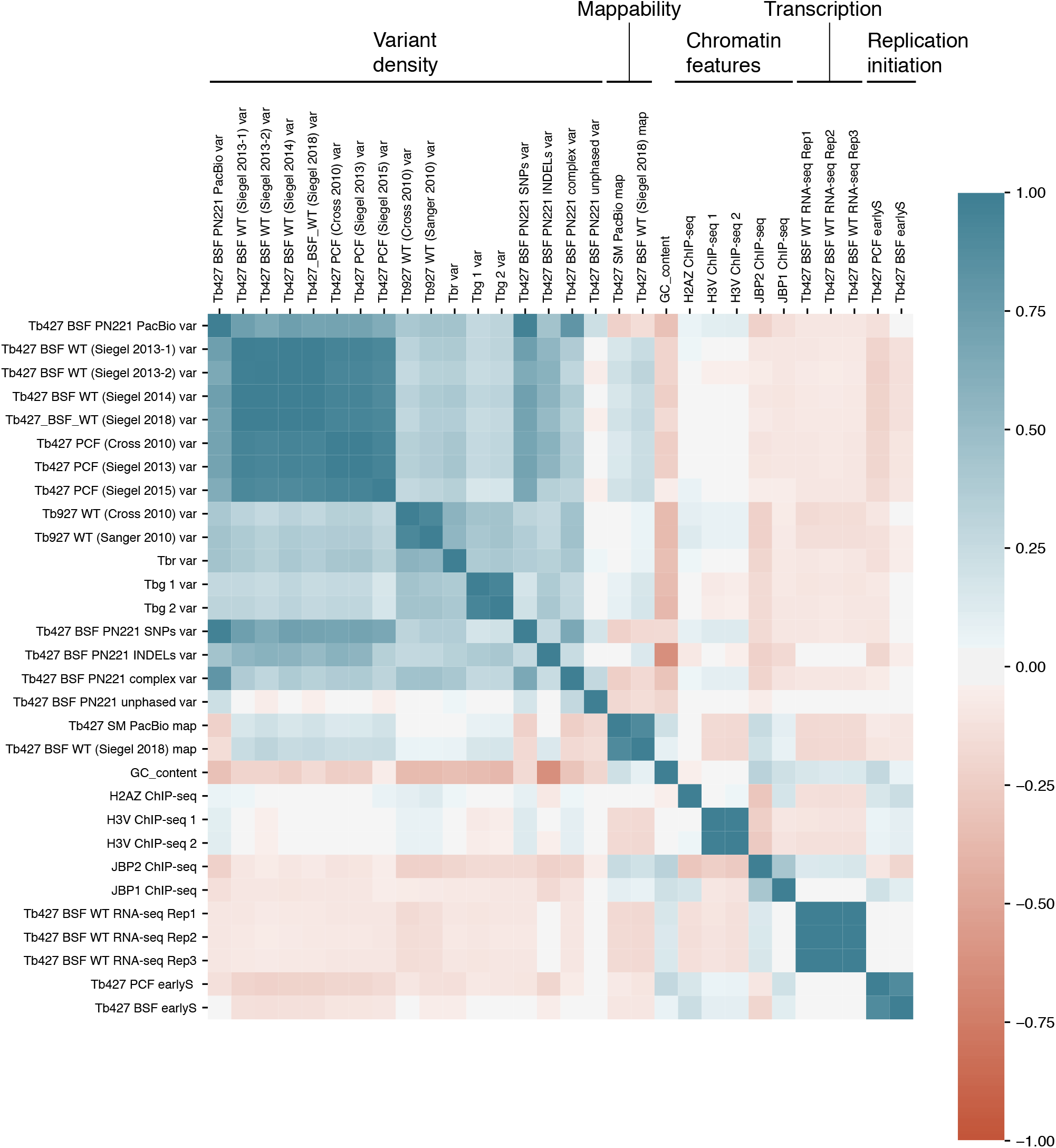
Correlation between variant density and different genomic features. Heatmap showing all vs all Pearson correlation for the variant density distribution of different *T. brucei* clones l, mappability, GC-content, enrichment of different chromatin factors, transcription and replication initiation. The description of the sequencing datasets used in this analysis, and their original publication, is listed in **Sup. Table S1**.

**Figure S3.**
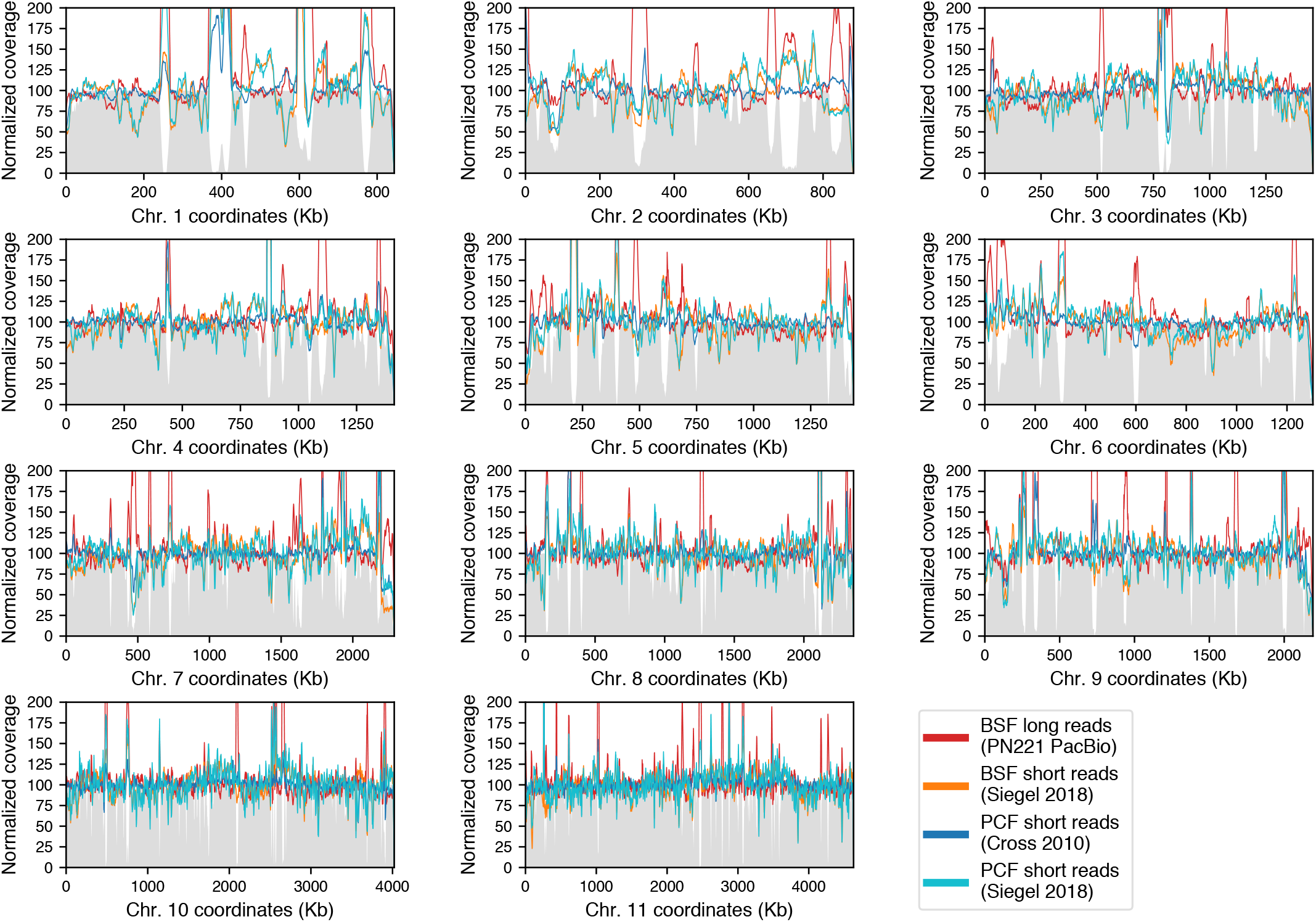
Genome-wide DNA-seq coverage in selected *T. brucei* Lister 427 clones. Normalized coverage along the eleven chromosomes for the DNA-seq datasets used to calculate variant density in Fig. 3. Mappability (in range 0-100) is shown as a grey filled-line.

**Figure S4.**
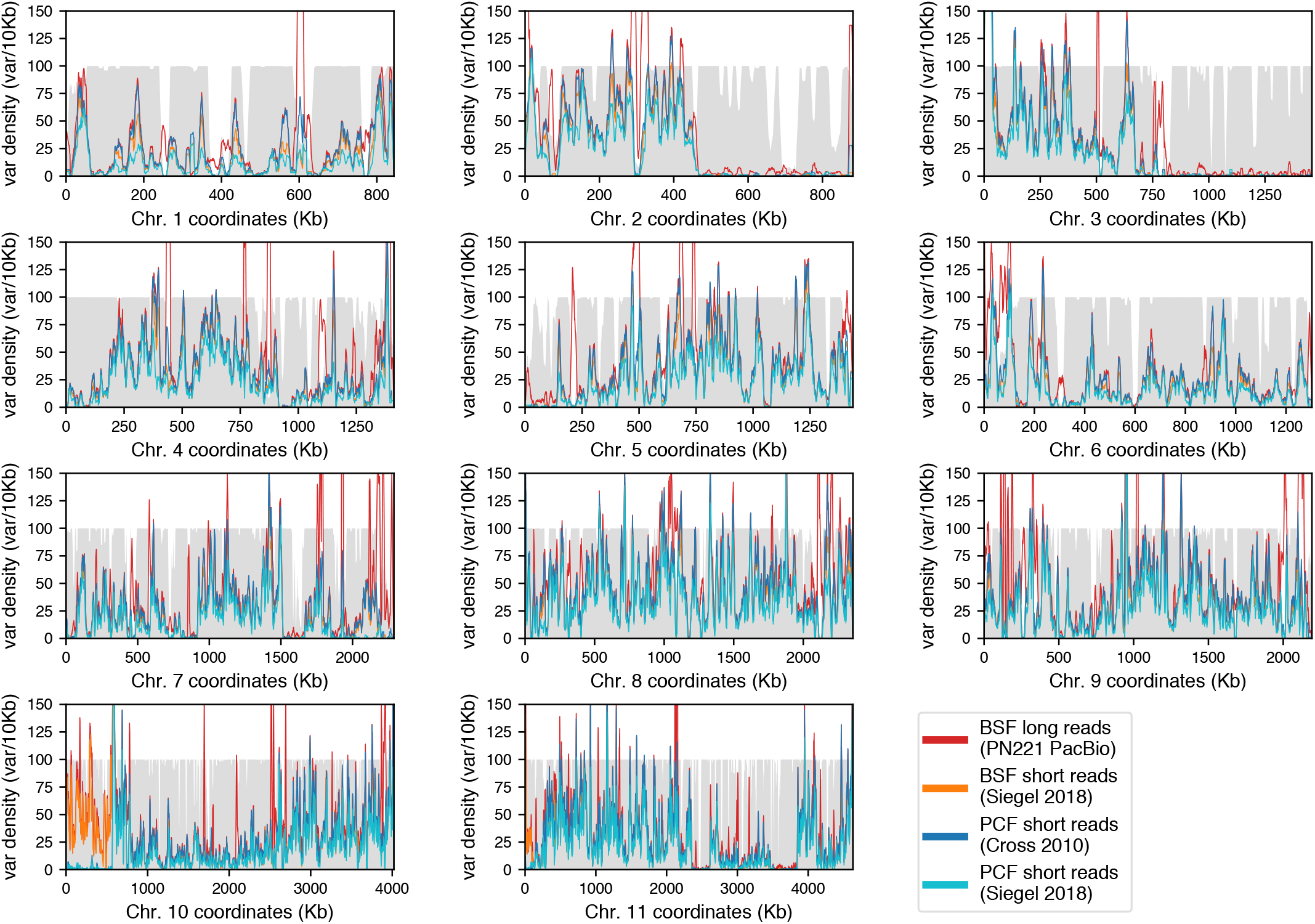
Genome-wide variant density in selected *T. brucei* Lister 427 clones. Variant density along the eleven chromosomes for the same DNA-seq dasets for which selected chromosomal regions were shown in Fig. 3. Mappability (in range 0-100) is shown as a grey filled-line.

**Figure S5.**
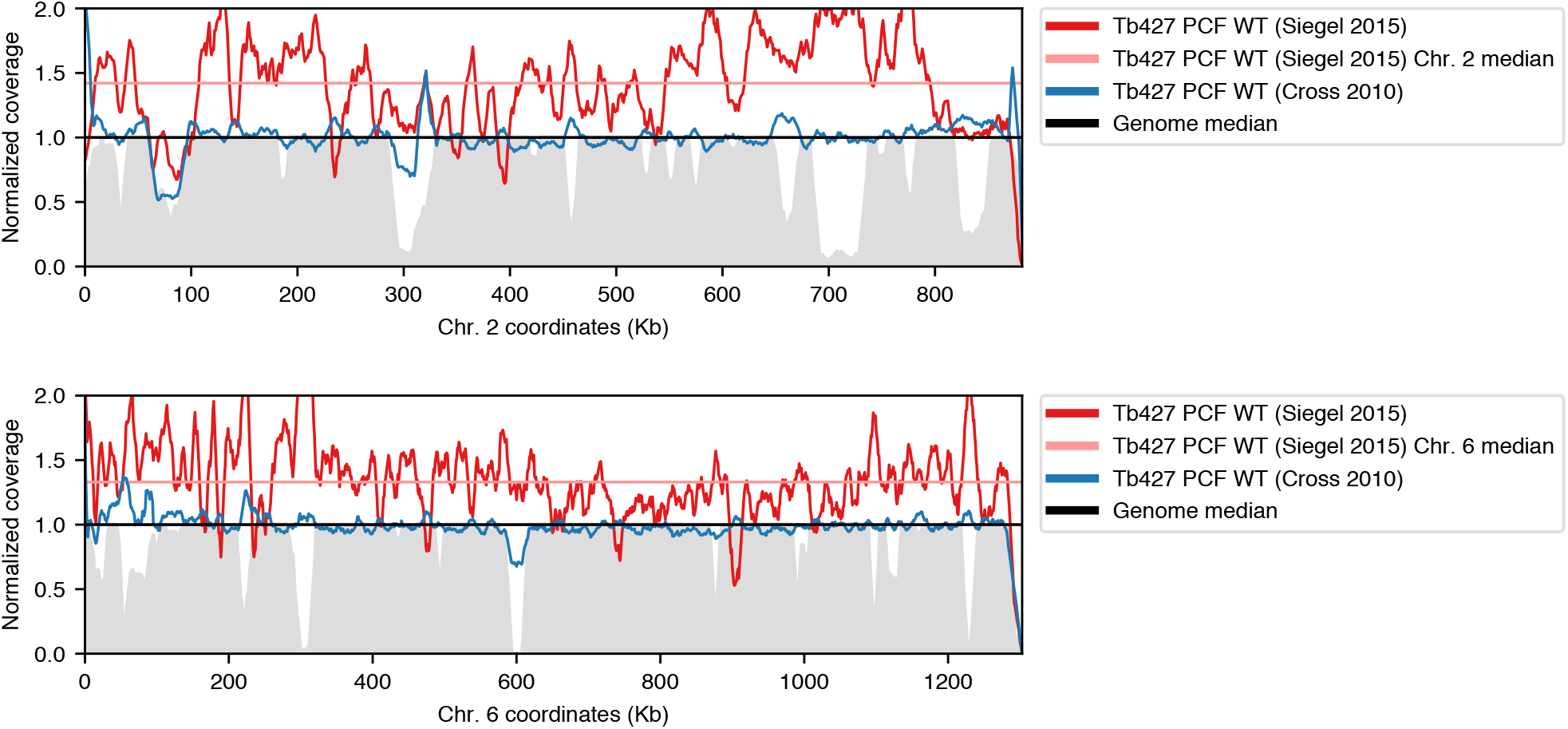
DNA-seq coverage in trisomic chromosomes in Tb427 PCF WT (Siegel 2015) clone. Normalized coverage density in chr 2 (top panel) and chr 6 (bottom panel) for Tb427 PCF WT (Siegel 2015) clone (dark red line) and its median (light red line), compared to Tb427 PCF WT (Cross 2010) clone (dark blue line). The genome median is set to 1 (straight black line). Mappability (in range 0-1) is shown as a grey filled-line.

**Figure S6.**
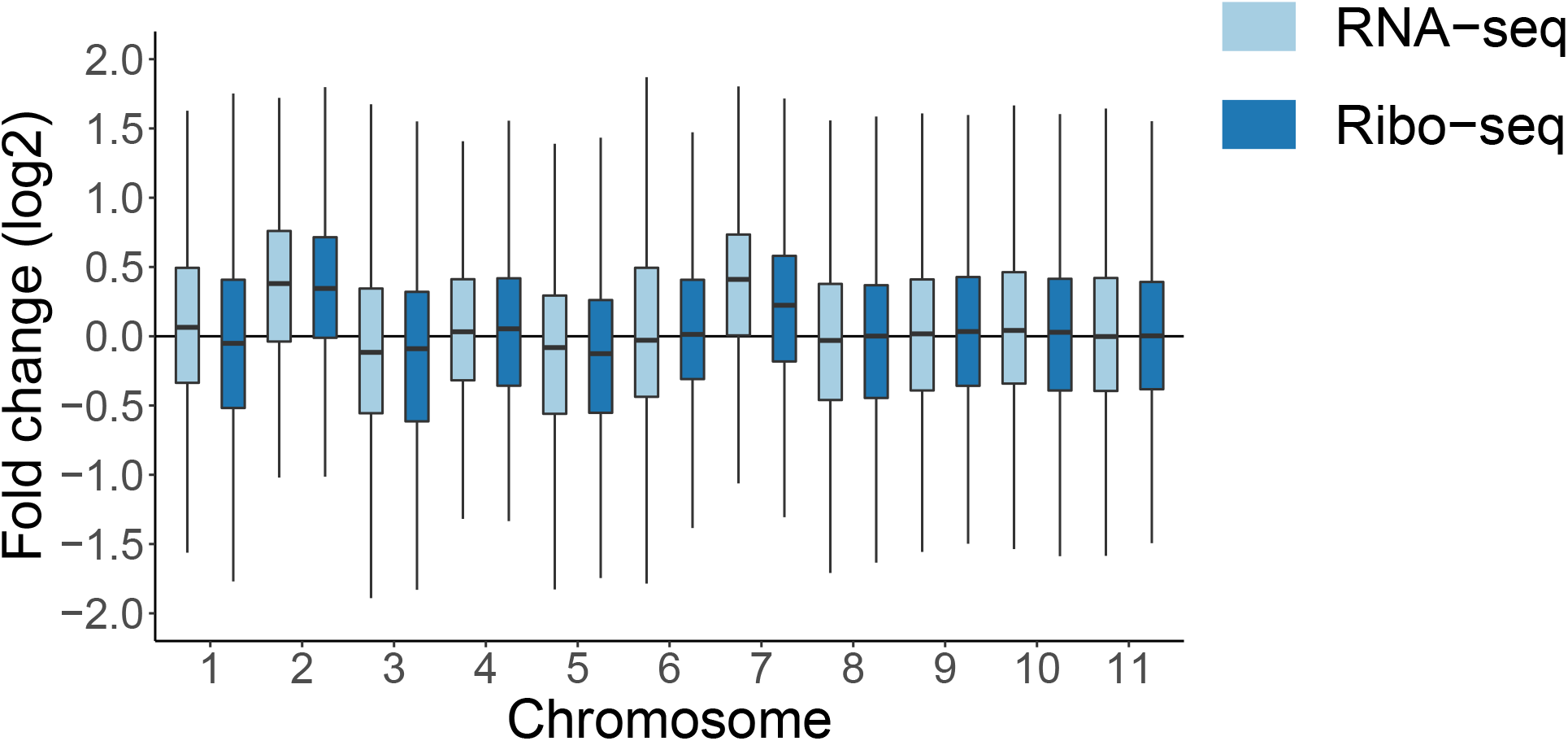
Genome-wide differential RNA-seq and Ribo-seq between aneuploid and diploid *T. brucei* clones. Log2-FoldChange RNA-seq (light blue) and Ribo-seq (dark-blue), pooled by chromosome, from a *T. brucei* Lister 427 PCF clone triploid for chr 2 and chr 7 over a diploid *T. brucei* TREU927 clone.

**Figure S7.**
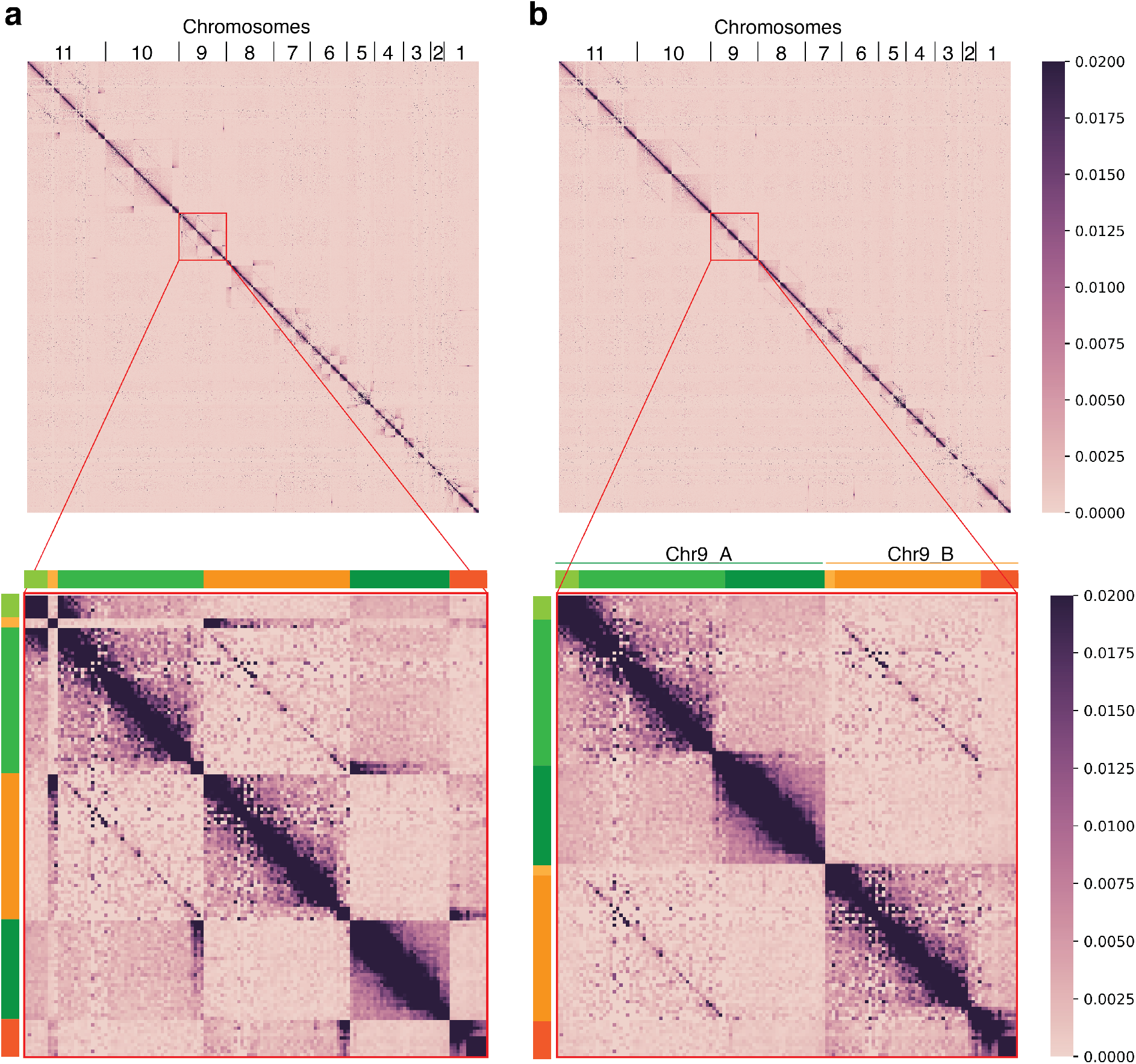
Hi-C interaction data between haploid-like subtelomers and phased-core alleles enables scaffolding of full homologous chromosomes. a) Hi-C interaction heatmap with the phased-core alleles and the haploid-like subtelomeres before scaffolding. Lower panel shows a zoom to the interactions between chr 9 phased-cores and haploid-like subtelomeres. b) The same as a) but after scaffolding the haploid-like subtelomeres to their interacting core allele.

**Figure S8.**
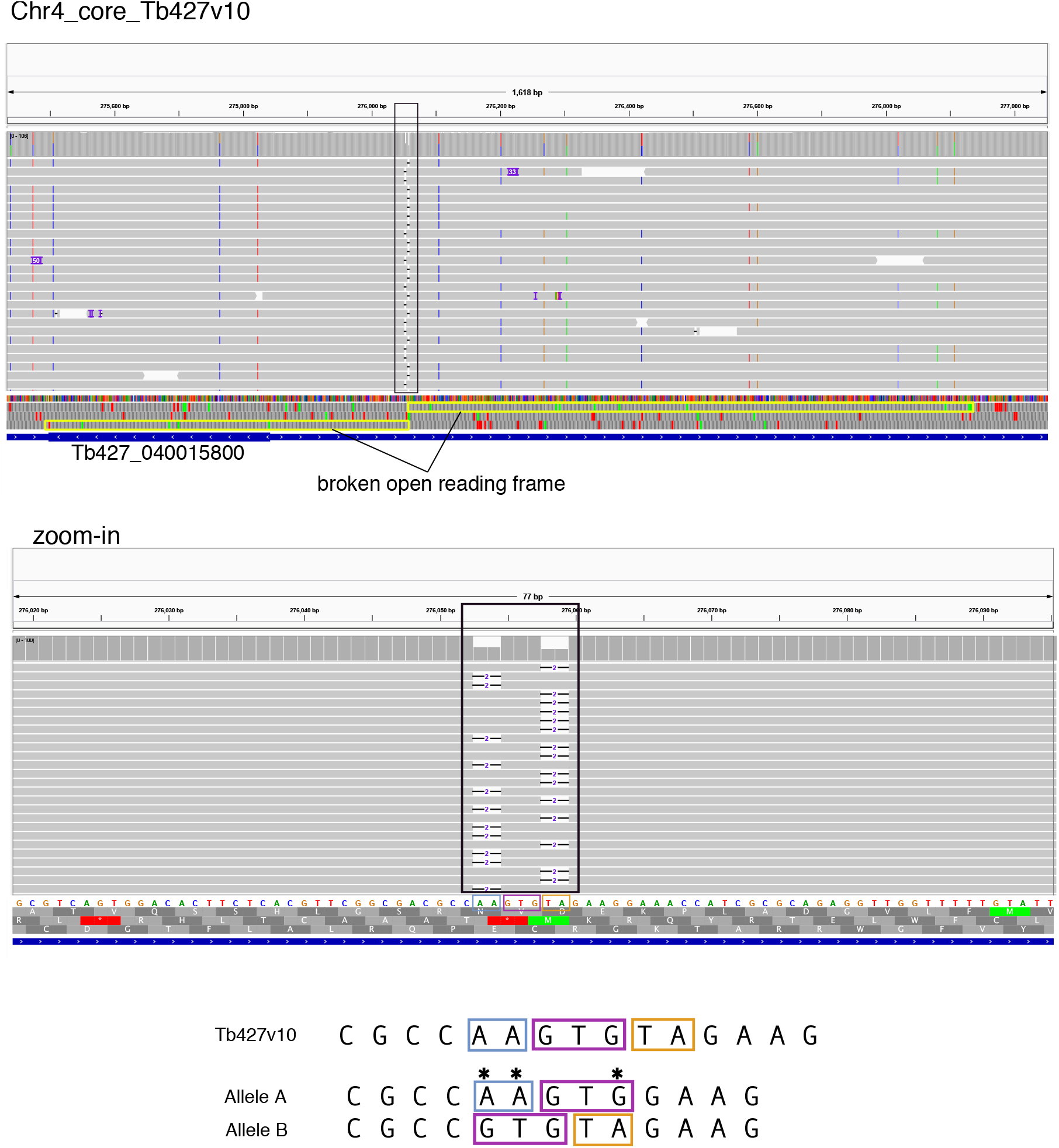
INDEL errors caused by nearby SNPs generating shifting sequence homology are fixed after allele-phasing. short error-corrected PacBio reads mapped to the collapsed Tb427v10 genome assembly. The upper panel is an Integrative Genome Viewer (IGV) screenshot showing a region from chr 4 containing the broken gene Tb427 040015800. Broken open reading frames are marked in yellow boxes in the translation panel. The conflicting region is marked with a black box. The lower IGV panel shows a zoom in to the conflicting region showing apparent deletions in different places for both haplotypes. At the bottom, the sequence present in Tb427v10 is shown, as well as the sequence in haplotype “A” and haplotype “B”, with violet boxes showing the shifted sequence homology between the alleles generated by the several heterozygote variants (marked with asterisks).

**Figure S9.**
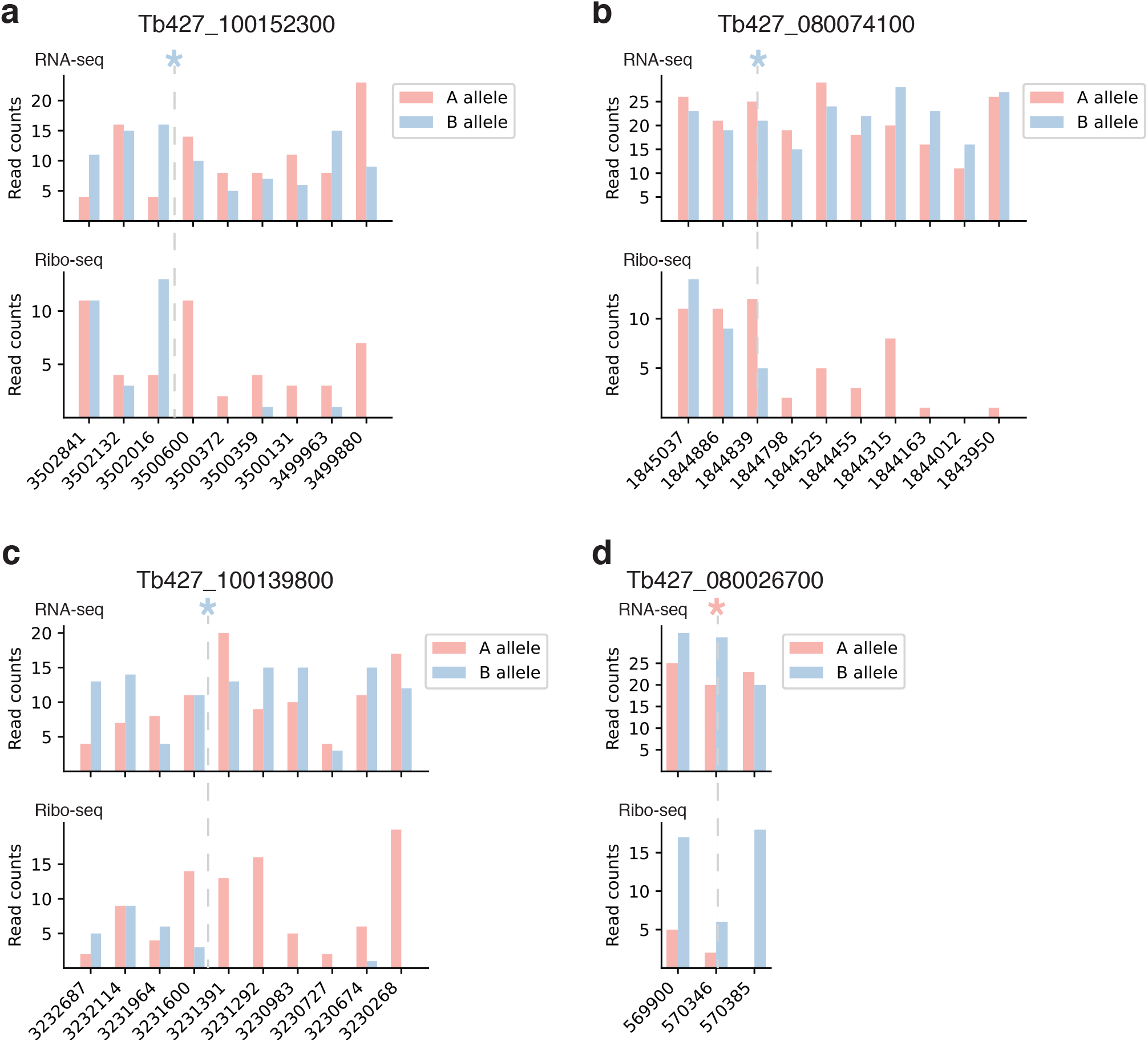
Allele-specific transcription and translation of genes with allele-specific premature termination codons. a-d) Examples of genes with allele-specific variants leading to a premature termination codon. The geneID of the genes in the Tb427v10 genome assembly is shown on top. Below, the barplots show RNA-seq and Ribo-seq read counts for both alleles in each of the variant position (chromosomal coordinates are indicated in the bottom label). An asterisk (and a grey dashed vertical line) indicates the position of the premature termination codon and the color of the asterisk indicates which allele has it.

